# Harmonised segmentation of neonatal brain MRI

**DOI:** 10.1101/2021.02.17.431611

**Authors:** Irina Grigorescu, Lucy Vanes, Alena Uus, Dafnis Batalle, Lucilio Cordero-Grande, Chiara Nosarti, A. David Edwards, Joseph V. Hajnal, Marc Modat, Maria Deprez

## Abstract

Deep learning based medical image segmentation has shown great potential in becoming a key part of the clinical analysis pipeline. However, many of these models rely on the assumption that the train and test data come from the same distribution. This means that such methods cannot guarantee high quality predictions when the source and target domains are dissimilar due to different acquisition protocols, or biases in patient cohorts. Recently, unsupervised domain adaptation (DA) techniques have shown great potential in alleviating this problem by minimizing the shift between the source and target distributions, without requiring the use of labelled data in the target domain. In this work, we aim to predict tissue segmentation maps on *T*_2_-weighted (*T*_2_w) magnetic resonance imaging (MRI) data of an unseen preterm-born neonatal population, which has both different acquisition parameters and population bias when compared to our training data. We achieve this by investigating two unsupervised DA techniques with the objective of finding the best solution for our problem. We compare the two methods with a baseline fully-supervised segmentation network and report our results in terms of Dice scores obtained on our ground truth test dataset. Moreover, we analyse tissue volumes and cortical thickness (CT) measures of the harmonised data on a subset of the population matched for gestational age (GA) at birth and postmenstrual age (PMA) at scan. Finally, we demonstrate the applicability of the harmonised cortical gray matter maps with an analysis comparing term and preterm-born neonates and a proof-of-principle investigation of the association between CT and a language outcome measure.

## 1 INTRODUCTION

Medical image deep learning has made incredible advances in solving a wide range of scientific problems, including tissue segmentation or image classification (Miotto et al., 2018). However, one major drawback of these methods is their applicability in a clinical setting, as many models rely on the assumption that the source and target domains are drawn from the same distribution. As a result, the efficiency of these models may drop drastically when applied to images which were acquired with acquisition protocols different than the ones used to train the models (Kamnitsas et al., 2017; Orbes-Arteaga et al., 2019).

A class of deep learning methods called DA techniques aims to address this issue by suppressing the domain shift between the training and test distributions. In general, DA approaches are either semi-supervised, which assume the existence of labels in the target dataset, or unsupervised, which assume the target dataset has no labels. For example, a common approach is to train a model on source domain images and fine-tune it on target domain data (Kushibar et al., 2019; Ghafoorian et al., 2017). Although these methods can give good results, they can become impractical as more often than not the existence of labels in the target dataset is limited or of poor quality. Unsupervised domain adaptation techniques (Ganin and Lempitsky, 2014; Kerfoot et al., 2019) offer a solution to this problem by minimizing the disparity between a source and a target domain, without requiring the use of labelled data in the target domain.

In our previous work (Grigorescu et al., 2020), we investigated two unsupervised DA methods with the aim of predicting brain tissue segmentations on 2D axial slices of *T*_2_ w MRI data of an unseen neonatal population. We proposed an additional loss term in one of the methods, in order to constrain the network to more realistic reconstructions. Our models were trained using as source domain a dataset with majority of term-born neonates and as target domain a preterm-only population acquired with a different protocol. We calculated mean cortical thickness measures for every subject in the two datasets and we performed an ANCOVA analysis in order to find group differences between the predicted source and target domains. This analysis showed that our proposed method achieved harmonisation of our two datasets in terms of cortical gray matter tissue segmentation maps. In this paper, we build on the aforementioned framework, which we expanded in three main ways. First, we build and train 3D neural networks in order to capture more information about the neonatal brain. Second, we extend the validation of our trained models to subsets of the two cohorts matched for GA and PMA, for which we analyse tissue volumes and global and local CT measures. Finally, we perform an analysis comparing term and preterm-born neonates on the harmonised cortical gray matter maps and we show the importance of harmonising the data by a proof-of-principle investigation of the association between cortical thickness and a language outcome measure.

## 2 MATERIAL AND METHODS

### 2.1 Data Acquisition and Preprocessing

The *T*_2_w MRI data used in this study was collected as part of two independent projects: the developing Human Connectome Project (dHCP^1^), and the Evaluation of Preterm Imaging (ePrime^2^) study. The dHCP data was acquired using a Philips 3T scanner and a 32-channels neonatal head coil (Hughes et al., 2017), using a *T*_2_w turbo spin echo (TSE) sequence with parameters: repetition time *T*_R_ = 12 s, echo time *T*_E_ = 156 ms, and overlapping slices with resolution 0.8 × 0.8 × 1.6 mm^3^. All data was motion corrected (Cordero-Grande et al., 2018; Kuklisova-Murgasova et al., 2012) and resampled to an isotropic voxel size of 0.5 mm^3^. The ePrime dataset was acquired with a Philips 3T system and an 8-channel phased array head coil, using a *T*_2_w fast-spin echo (FSE) sequence with parameters: repetition time *T*_R_ = 14.73 s and echo time *T*_E_ = 160 ms (Ball et al., 2017). Images were acquired with a voxel size of 0.86 × 0.86 × 2 mm, with 1 mm overlap.

Our two datasets comprise of 402 MRI scans of infants born between 23 — 42 weeks GA at birth and scanned at term-equivalent age (after 37 weeks PMA) as part of the dHCP pipeline, and a dataset of 485 MRI scans of infants born between 23 — 33 weeks GA and scanned at term-equivalent age as part of the ePrime project. Figure 1 shows their age distribution.

**Figure 1.**
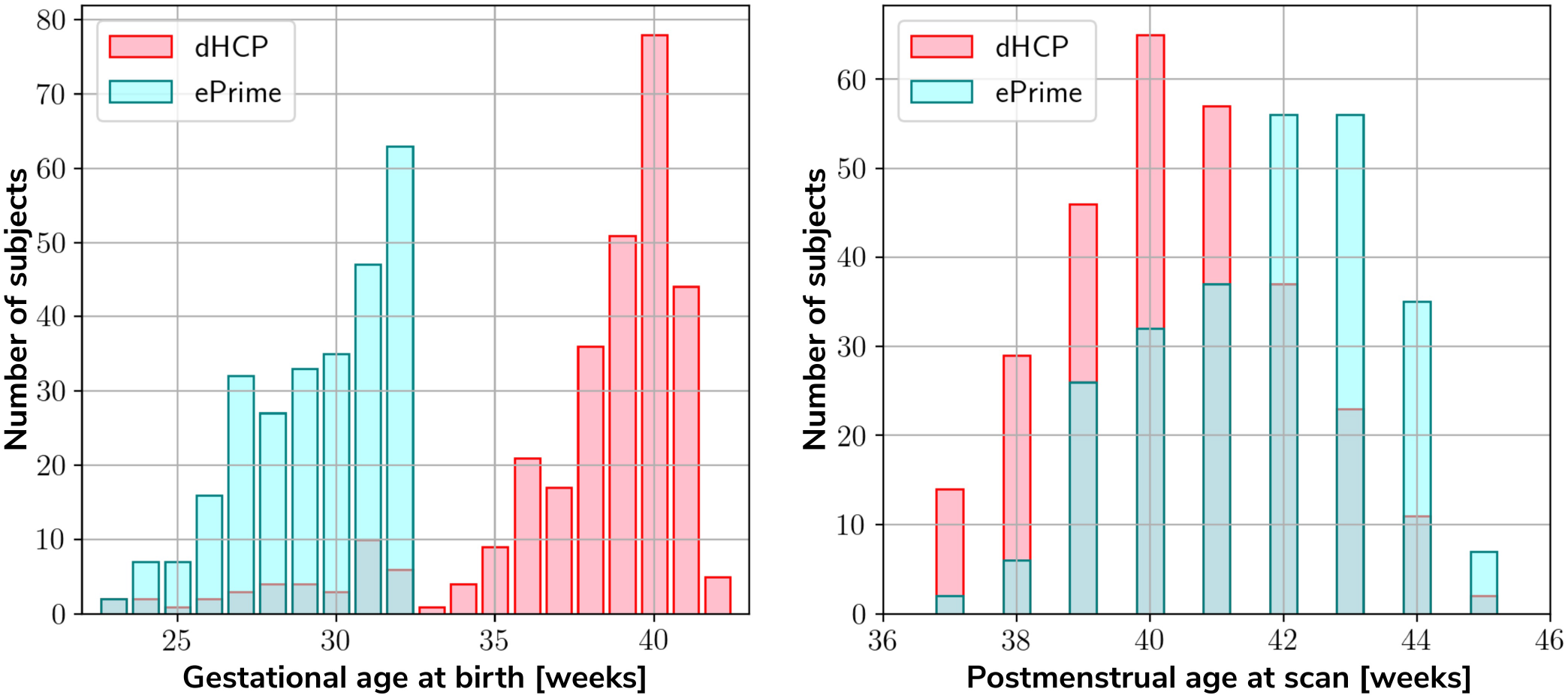
Age distribution of the subjects in our datasets, showing both their GA at birth, as well as their PMA at scan.

Both datasets were pre-processed prior to being used by the deep learning algorithms. The ePrime volumes were linearly upsampled to 0.5 mm^3^ isotropic resolution to match the resolution of our source (dHCP) dataset. Both dHCP and ePrime datasets were rigidly aligned to a common 40 weeks gestational age atlas space (Schuh et al., 2018) using the MIRTK (Rueckert et al., 1999) software toolbox. Then, skull-stripping was performed on all of our data using the brain masks obtained with the Draw-EM pipeline for automatic brain MRI segmentation of the developing neonatal brain (Makropoulos et al., 2018). Ground truth tissue segmentation maps were obtained using the same pipeline (Draw-EM) for both cohorts.

To train our networks, we split our datasets into 80% training, 10% validation and 10% test (see Table 1), keeping the distribution of ages at scan as close to the original as possible. We used the validation sets to keep track of our models’ performance during training, and the test sets to report our final models’ results and showcase their capability to generalize.

**Table 1.**
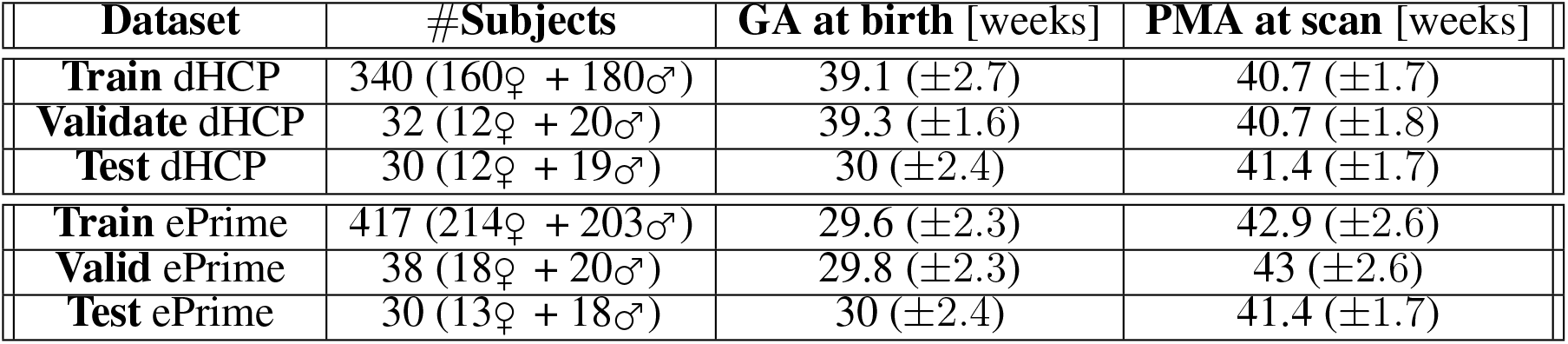
Number of scans in different datasets used for training, validation and testing the models, together with their mean GA and PMA

| Dataset | #Subjects | GA at birth [weeks] | PMA at scan [weeks] |
| --- | --- | --- | --- |
| **Train** dHCP | 340 (160♀ + 180♂) | 39.1 ($\pm 2.7$) | 40.7 ($\pm 1.7$) |
| **Validate** dHCP | 32 (12♀ + 20♂) | 39.3 ($\pm 1.6$) | 40.7 ($\pm 1.8$) |
| **Test** dHCP | 30 (12♀ + 19♂) | 30 ($\pm 2.4$) | 41.4 ($\pm 1.7$) |
| **Train** ePrime | 417 (214♀ + 203♂) | 29.6 ($\pm 2.3$) | 42.9 ($\pm 2.6$) |
| **Valid** ePrime | 38 (18♀ + 20♂) | 29.8 ($\pm 2.3$) | 43 ($\pm 2.6$) |
| **Test** ePrime | 30 (13♀ + 18♂) | 30 ($\pm 2.4$) | 41.4 ($\pm 1.7$) |

### 2.2 Unsupervised domain adaptation models

To investigate the best solution for segmenting our target dataset (ePrime), we compared three independently trained deep learning models:

- **Baseline**. A 3D U-Net (Ronneberger et al., 2015) trained on the source dataset (dHCP) only and used as a baseline segmentation network (see Figure 2).
- **Adversarial domain adaptation in the latent space**. A 3D U-Net segmentation network trained on source (dHCP) volumes, coupled with a discriminator trained on both source (dHCP) and target (ePrime) datasets (see Figure 3). This solution is similar to the one proposed by (Kamnitsas et al., 2017) where the aim was to train the segmentation network such that it becomes agnostic to the data domain.
- **Adversarial domain adaptation in the image space**. Two 3D U-Nets, one acting as a generator, and a second one acting as a segmentation network, coupled with a discriminator trained on both real and fake ePrime volumes. The segmentation network is trained to produce tissue maps of the fake ePrime-like volumes created by the generator (see Figure 4). The normalised cross correlation (NCC) loss is added to the generator network to enforce image similarity between real and synthesised images, a solution which was previously proposed by (Grigorescu et al., 2020).

**Figure 2.**
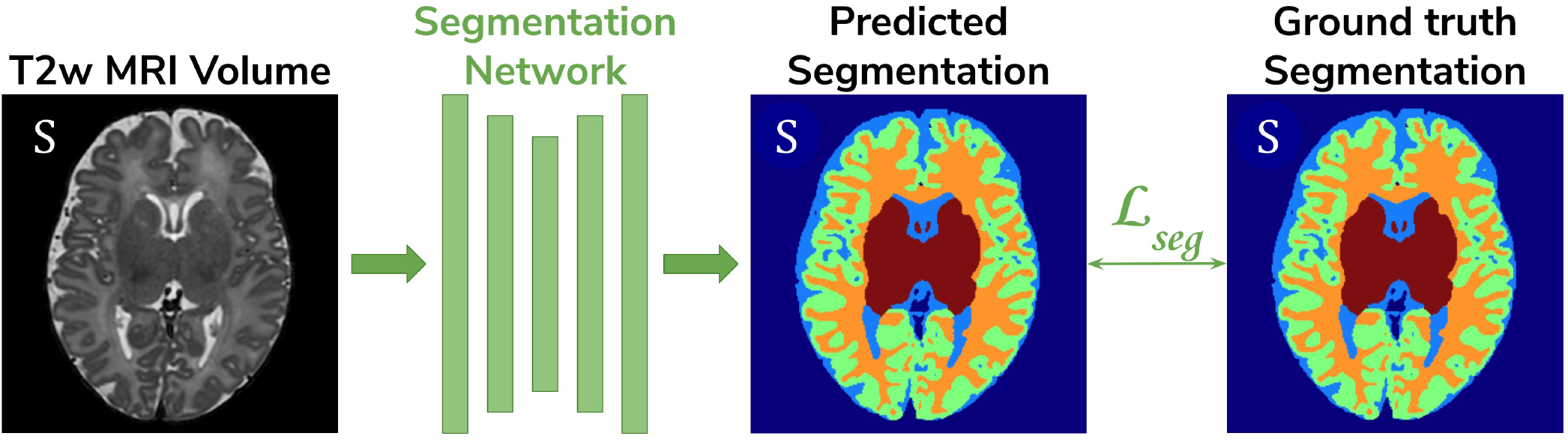
The baseline model consists of a 3D U-Net trained to segment source (dHCP) volumes. The input *T*_2_w MRI images, the predicted segmentation and the ground truth segmentations are marked with S as they all belong to the source (dHCP) dataset.

**Figure 3.**
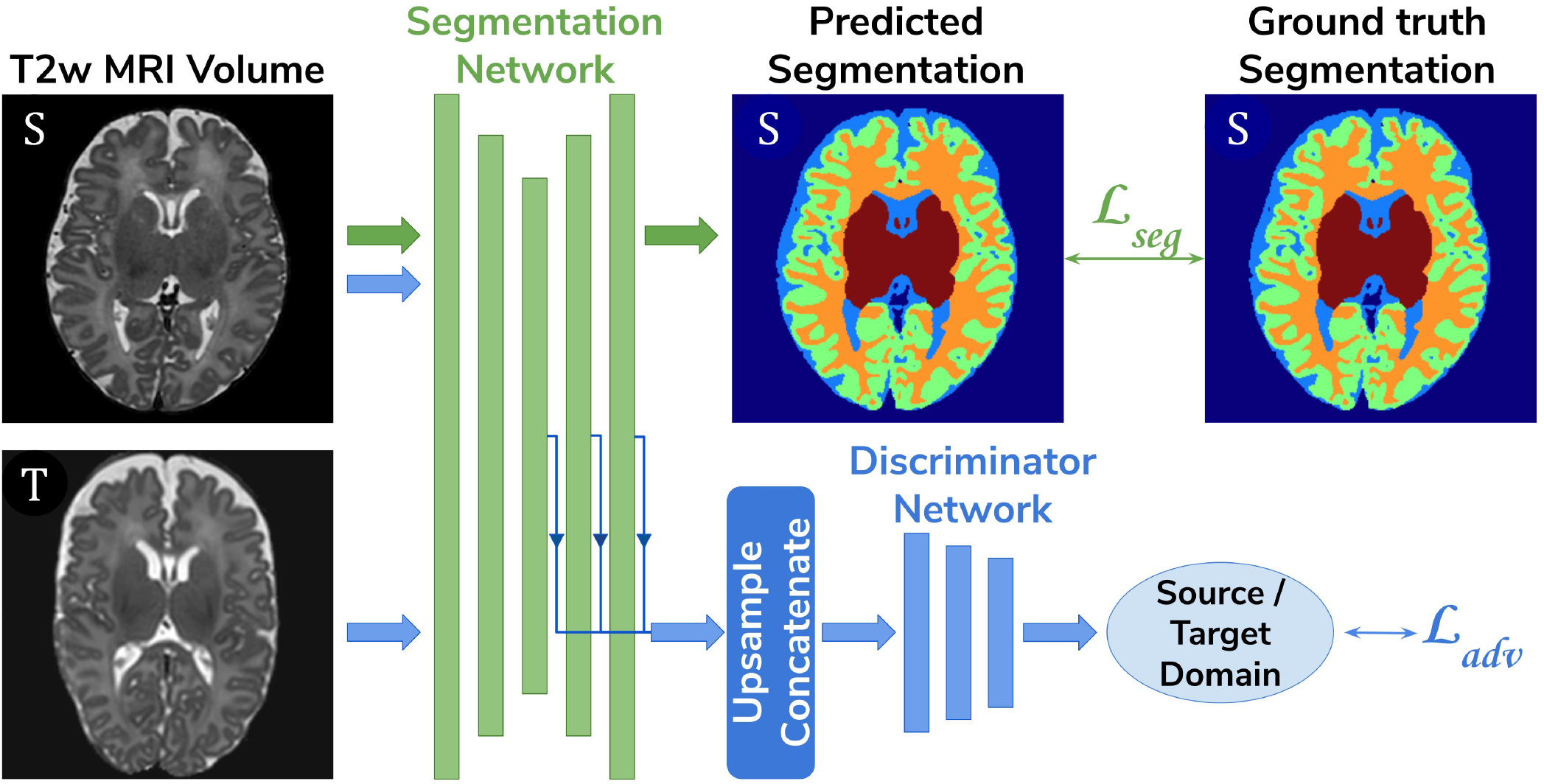
The latent space domain adaptation setup consists of a 3D U-Net trained to segment the source (dHCP) *T*_2_w MRI volumes, coupled with a discriminator network which forces the segmentation network to learn domain-invariant features. Both source (dHCP) and target (ePrime) images are fed to the segmentation network, but only source (dHCP) ground truth labels are used to compute the segmentation loss. Source domain images are marked with S, while target domain images are marked with T, respectively.

**Figure 4.**
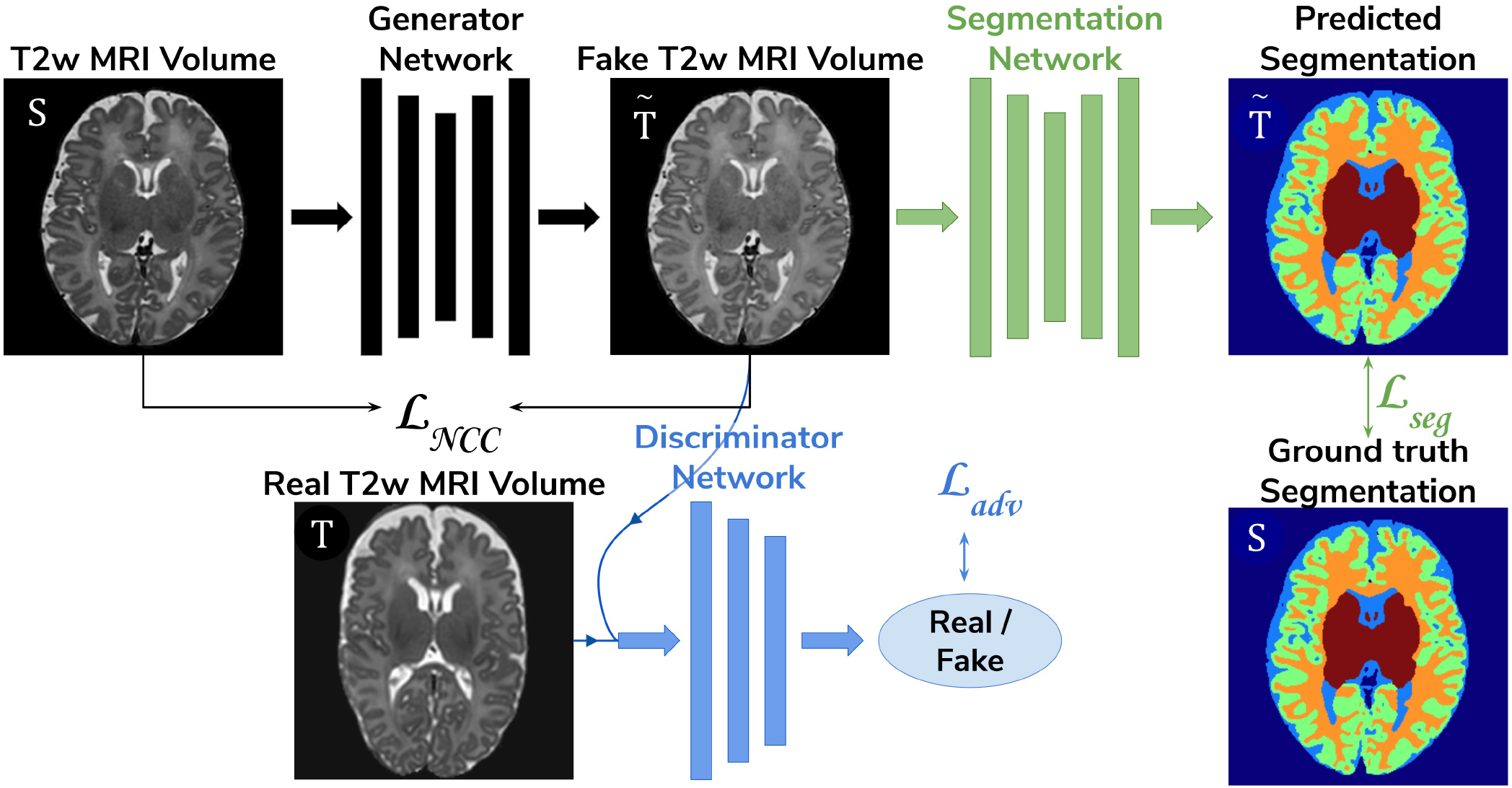
The image space domain adaptation setup uses a generator network to produce ePrime-like *T*_2_w MRI images (marked with 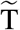), which are then used as input into the segmentation network. The discriminator is trained to distinguish between real (ePrime) and fake (ePrime-like) volumes, while the generator is trained to produce realistic images in order to fool the discriminator. The NCC loss enforces image similarity between real and synthesised volumes.

To further validate the harmonised tissue maps, we trained an additional network (a 3D U-Net) to segment binary cortical tissue maps into 11 cortical substructures (see Table 2) based on anatomical groupings of cortical regions derived from the Draw-EM pipeline. The key reasons for training an extra network are: first, we avoid the time consuming task of label propagation between our available dHCP ground truth segmentations and predicted ePrime maps, and second, we can train this network using ground truth cortical segmentations, and apply it on any brain cortical gray matter maps as in this case there will be no intensity shift between target and source distributions.

**Table 2.**
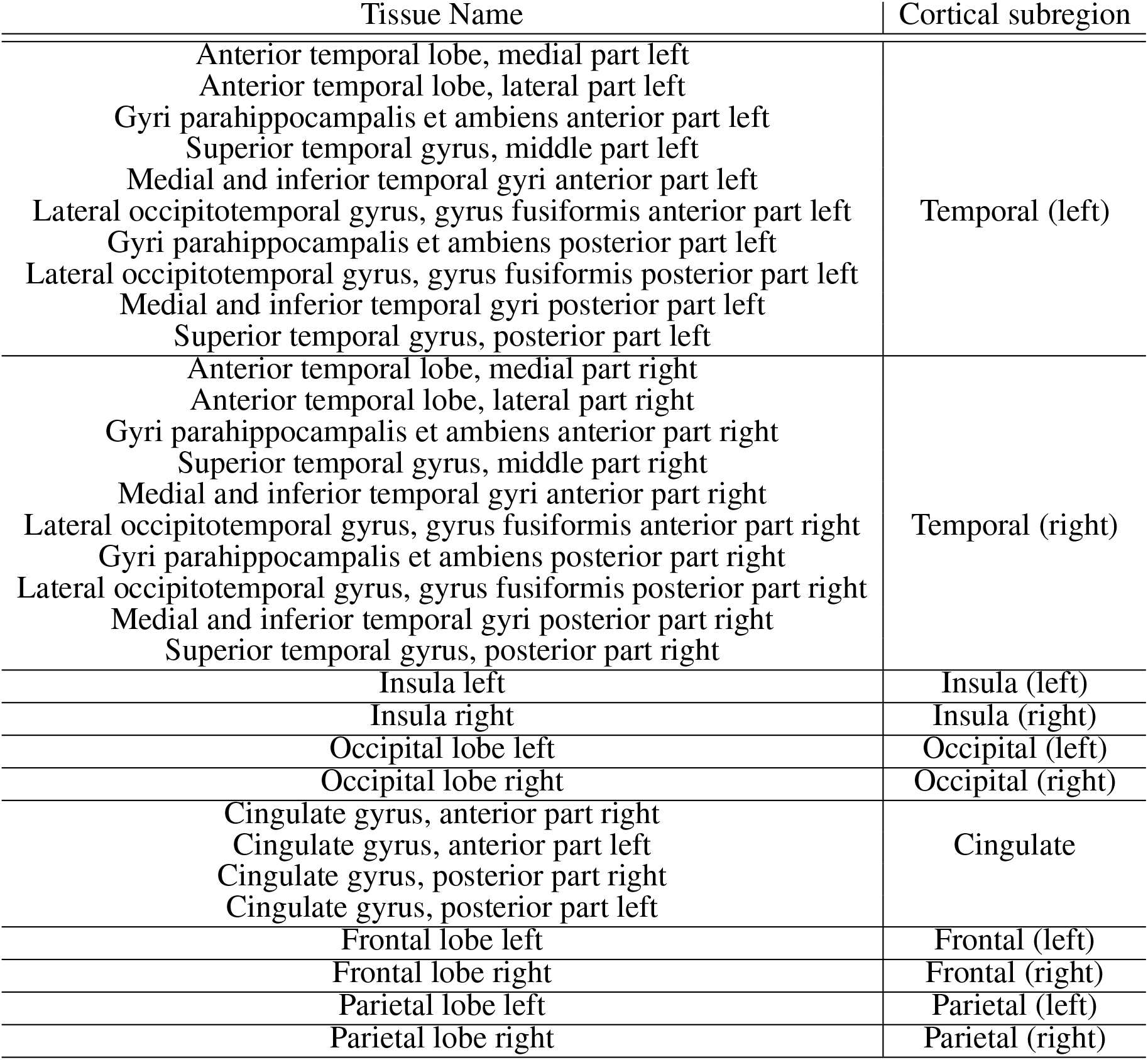
Grouping of cortical substructures showing their original tissue name obtained from Draw-EM (Makropoulos et al., 2018) on the first column and their corresponding cortical subregion on the second column.

| Tissue Name | Cortical subregion |
| --- | --- |
| Anterior temporal lobe, medial part left<br>Anterior temporal lobe, lateral part left<br>Gyri parahippocampalis et ambiens anterior part left<br>Superior temporal gyrus, middle part left<br>Medial and inferior temporal gyri anterior part left<br>Lateral occipitotemporal gyrus, gyrus fusiformis anterior part left<br>Gyri parahippocampalis et ambiens posterior part left<br>Lateral occipitotemporal gyrus, gyrus fusiformis posterior part left<br>Medial and inferior temporal gyri posterior part left<br>Superior temporal gyrus, posterior part left | Temporal (left) |
| Anterior temporal lobe, medial part right<br>Anterior temporal lobe, lateral part right<br>Gyri parahippocampalis et ambiens anterior part right<br>Superior temporal gyrus, middle part right<br>Medial and inferior temporal gyri anterior part right<br>Lateral occipitotemporal gyrus, gyrus fusiformis anterior part right<br>Gyri parahippocampalis et ambiens posterior part right<br>Lateral occipitotemporal gyrus, gyrus fusiformis posterior part right<br>Medial and inferior temporal gyri posterior part right<br>Superior temporal gyrus, posterior part right | Temporal (right) |
| Insula left | Insula (left) |
| Insula right | Insula (right) |
| Occipital lobe left | Occipital (left) |
| Occipital lobe right | Occipital (right) |
| Cingulate gyrus, anterior part right<br>Cingulate gyrus, anterior part left<br>Cingulate gyrus, posterior part right<br>Cingulate gyrus, posterior part left | Cingulate |
| Frontal lobe left | Frontal (left) |
| Frontal lobe right | Frontal (right) |
| Parietal lobe left | Parietal (left) |
| Parietal lobe right | Parietal (right) |

### 2.3 Network Architectures

The segmentation networks in all three setups and the generator used in the adversarial domain adaptation in the image space model have the same architecture, consisting of 5 encoding-decoding branches with 16, 32, 64, 128 and 256 channels, respectively. The encoder blocks use 3^3^ convolutions (with a stride of 1), instance normalisation (Ulyanov et al., 2016) and LeakyReLU activations. A 2^3^ average pooling layer is used after the first down-sampling block, while the others use 2^3^ max pooling layers. The decoder blocks consist of 3^3^ convolutions (with a stride of 1), instance normalisation (Ulyanov et al., 2016), LeakyReLU activations, and, additionally, 3^3^ transposed convolutions. The segmentation network outputs a 7-channel 3D volume (of the same size as the input image), corresponding to our 7 classes: background, cerebrospinal fluid (CSF), cortical gray matter (cGM), white matter (WM), deep gray matter (dGM), cerebellum and brainstem. The generator network’s last convolutional layer is followed by a Tanh activation and outputs a single channel image.

For our unsupervised domain adaptation models (Figures 3 and 4) we used a PatchGAN discriminator as proposed in (Isola et al., 2016). Its architecture consists of 4 layers of 3D convolutions (of 64, 128, 256 and 512 channels, respectively), instance normalisation and LeakyReLU activations.

The cortical parcellation network has the same architecture as the tissue segmentation network, but outputs a 12-channel 3D volume corresponding to the following cortical substructures: frontal left, frontal right, cingulate, temporal left, temporal right, insula left, insula right, parietal left, parietal right, occipital left, and occipital right, respectively. The last class represents the background.

### 2.4 Training

The baseline segmentation network (Figure 2) was trained by minimizing the generalised Dice loss (Sudre et al., 2017) between the predicted and the ground truth segmentation maps (Equation 1).

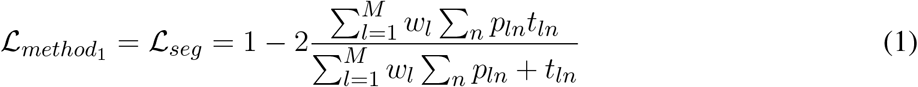

where 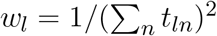 is the weight of the l*^th^* tissue type, *p_ln_* is the predicted probabilistic map of the l*^th^* tissue type at voxel n, t_ln_ is the target label map of the l^th^ tissue type at voxel n, and M is the number of tissue classes. While training, we used the Adam optimizer with its default parameters and a decaying cyclical learning rate scheduler (Smith, 2015) with a base learning rate of 2 · 10^−6^ and a maximum learning rate of 2 · 10^−3^.

The segmentation network from the adversarial domain adaptation in the latent space model was trained to produce tissue maps on the source (dHCP) volumes. In addition, both target and source volumes were fed to the segmentation network, while the feature maps obtained from every level of its decoder arm were passed to the discriminator network which acted as a domain classifier. This was done after either up-sampling or down-sampling the feature maps to match the volume size of the second deepest layer. This model was trained by minimizing a Cross-Entropy loss between predicted and assigned target labels representing our two domains. The final loss function for our second model was therefore made up of the generalised Dice loss and an adversarial loss:

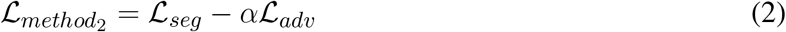

where *α* was a hyperparameter increased linearly from 0 to 0.05 starting at epoch 20, and which remained equal to 0.05 from epoch 50 onward. As explained in (Kamnitsas et al., 2017), the aim was to both maximise the domain classification loss, while minimizing the segmentation loss. The segmentation network was trained similarly to the baseline model, while the discriminator network was trained using the Adam optimiser with *β_1_* = 0.5 and *β_2_* = 0.999, and a linearly decaying learning rate scheduler starting from 2 · 10 ^−3^.

The generator network used in the image space domain adaptation approach was trained to produce fake ePrime volumes, while the segmentation network was trained using the same loss function, optimizer and learning rate scheduler as in the other two methods. For both the discriminator and the generator networks the Adam optimizer with *β*_1_ = 0.5 and *β*_2_ = 0.999 was used, together with a linearly decaying learning rate scheduler starting from 2 · 10^−3^. The loss function of the discriminator was similar to that of the Least Squares GAN (Mao et al., 2016): 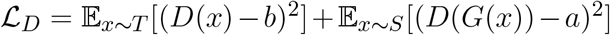 where *a* signified the label for fake volumes and *b* was the label for real volumes. The generator and the segmentation network are trained together using the following loss:

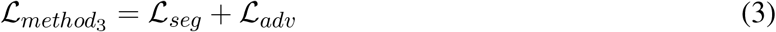

where 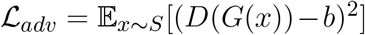. An additional NCC loss was used between the real and the generated volumes in order to constrain the generator to produce realistic looking ePrime-like images. Without the additional NCC loss, the generator tends to produce synthesized images with an enlarged CSF boundary in order to match the preterm-only born distribution found in the ePrime dataset, as we previously shown in (Grigorescu et al., 2020).

These three methods were trained with and without data augmentation for 100 epochs, during which we used the validation sets to inform us about our models’ performance and to decide on the best performing models. For data augmentation we applied: random affine transformations (with rotation angles 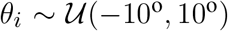 and/or scaling values 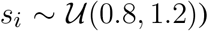, random motion artefacts (corresponding to rotations of 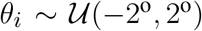 and translations of 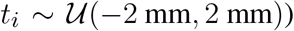, and random MRI spike and bias field artifacts (Perez-García et al., 2020). The cortical parcellation network was trained in a similar fashion as the baseline tissue segmentation network, with data augmentation in the form of random affine transformations (with the same parameters as above).

The test set was used to report our final models’ results and to showcase their capability to generalize on the source domain. Finally, we produced tissue segmentation maps for all the subjects in our datasets, and used them as input into ANT’s DiReCT algorithm (Tustison et al., 2013) to compute cortical thickness measures. To validate our results, we compared cortical thickness measures between subsets of the two cohorts matched for GA and PMA, for which we expect no significant difference in cortical thickness if the harmonisation was successful. We also assessed the association between PMA and cortical thickness in the two cohorts.

## 3 RESULTS

### 3.1 dHCP test dataset

#### Baseline and domain adaptation models

Figure 5 summarizes the results of our trained models when applied on the test dataset of the source domain (dHCP) for which we have ground truth segmentations. The figure shows the mean Dice scores computed between the predicted tissue segmentation maps and the ground truth labels for each of the six trained models. The model that obtained the highest mean Dice scores is highlighted with the yellow diamond. Out of the six models, the *baseline with augmentation* and *image with augmentation* methods performed best on the source domain test dataset for CSF, dGM, cerebellum and brainstem, with no significant difference between them. For cGM and WM, the best performance was obtained by the *baseline with augmentation* model, while the domain adaptation methods showed a slight decrease in performance. The three models trained without augmentation always performed significantly worse than their augmented counterparts. These results show that our trained models were able to generalise to unseen source domain data, and that the performance on the dHCP dataset was not compromised by using domain adaption techniques.

**Figure 5.**
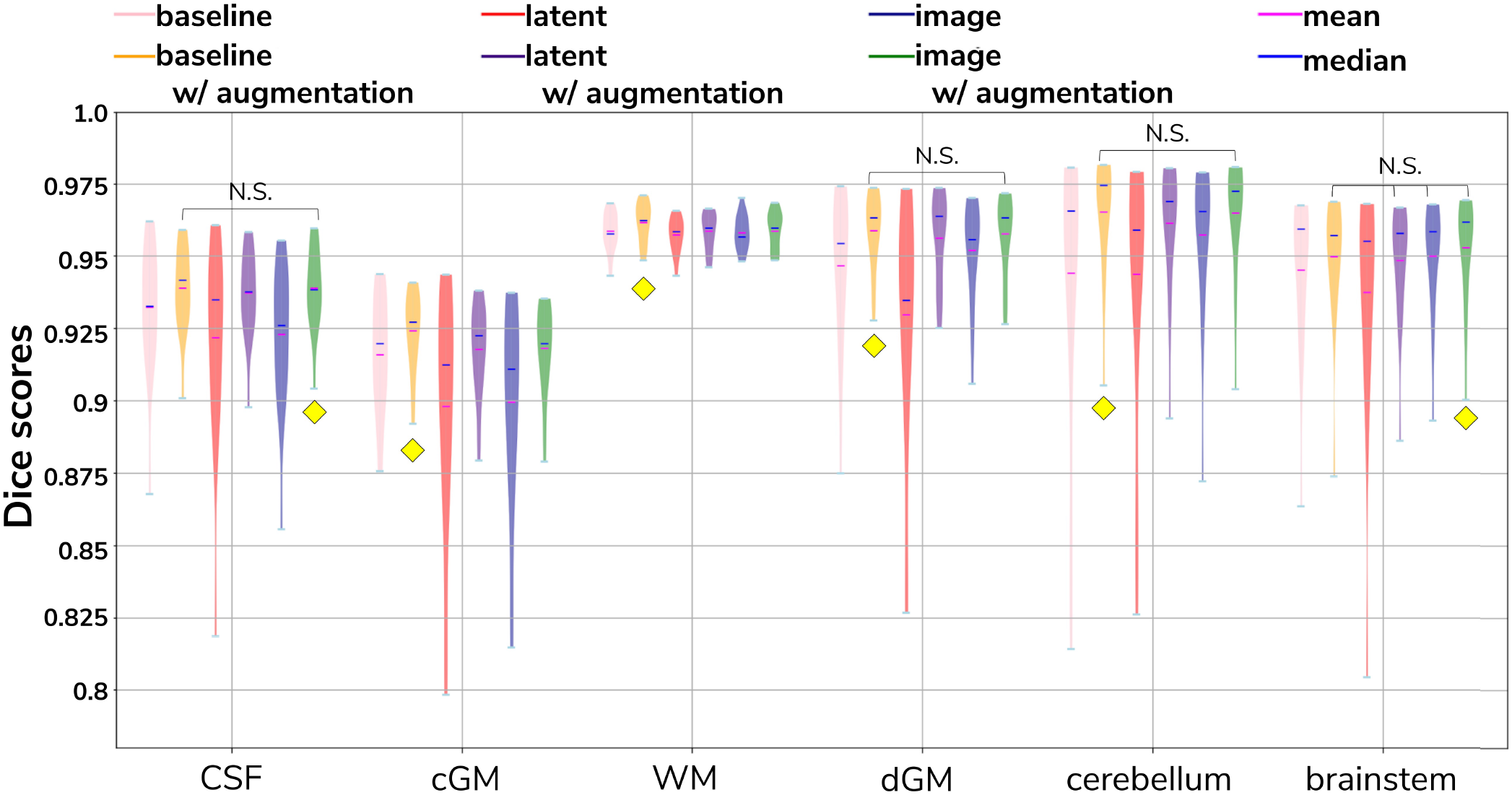
The results on our dHCP test dataset for all six methods. Models with non-significant differences in mean Dice Scores when compared to the *baseline with augmentation* method are shown above each pair. The yellow diamond highlights the model which obtained the highest mean Dice score for its respective tissue type.

#### Cortical parcellation network

Table 3 summarizes the results of applying the trained cortical parcellation network on the dHCP test dataset. When compared with the ground truth segmentations obtained using the Draw-EM pipeline (Makropoulos et al., 2018), the network obtained an overall mean Dice score of 0.97.

**Table 3.** Dice Scores obtained on the dHCP test set for the trained cortical parcellation network.

| Tissue | min | max | mean | Tissue | min | max | mean |
| --- | --- | --- | --- | --- | --- | --- | --- |
| Frontal (left) | 0.98 | 0.99 | 0.99 | Frontal (right) | 0.98 | 0.99 | 0.99 |
| Temporal (left) | 0.96 | 0.99 | 0.98 | Temporal (right) | 0.97 | 0.98 | 0.98 |
| Insula (left) | 0.95 | 0.97 | 0.96 | Insula (right) | 0.95 | 0.97 | 0.96 |
| Parietal (left) | 0.96 | 0.98 | 0.97 | Parietal (right) | 0.96 | 0.98 | 0.97 |
| Occipital (left) | 0.94 | 0.98 | 0.97 | Occipital (right) | 0.95 | 0.98 | 0.97 |
| Cingulate | 0.93 | 0.97 | 0.96 |  |  |  |  |

### 3.2 Validation of data harmonisation

In order to evaluate the extent to which each of the trained models managed to harmonise the segmentation maps of the two cohorts, we looked at tissue volumes and mean cortical thickness measures between subsamples of the dHCP (*N* = 30; median GA = 30.50 weeks; median PMA = 41.29 weeks) and ePrime (*N* = 30; median GA = 30.64 weeks; median PMA = 41.29 weeks) cohort which showed comparable GA at birth and PMA at time of scan (see Table 1). For these two cohort subsamples with similar GA and PMA, we expected both volumes and cortical thickness measures not to differ after applying the harmonisation procedures. We also investigated the relationship between PMA and volumes and cortical thickness respectively, before and after applying the harmonisation. Linear regressions were performed in the comparable data subsets testing the effects of PMA and cohort on volumes (or cortical thickness), controlling for GA and sex.

#### Volumes

Figure 6 shows the tissue volumes for both the original and the predicted segmentations. Significant volume differences between the two subsamples (i.e., significant effect of cohort in the regression model) are reported above each tested model. To summarise, the *image with augmentation* model performed best, by showing no significant differences in the two cohorts for cortical gray matter, white matter, deep gray matter, cerebellum and brainstem. The cerebrospinal fluid volumes were significantly different between the two cohorts for all our trained models, as well as for the original ePrime segmentation masks.

**Figure 6.**
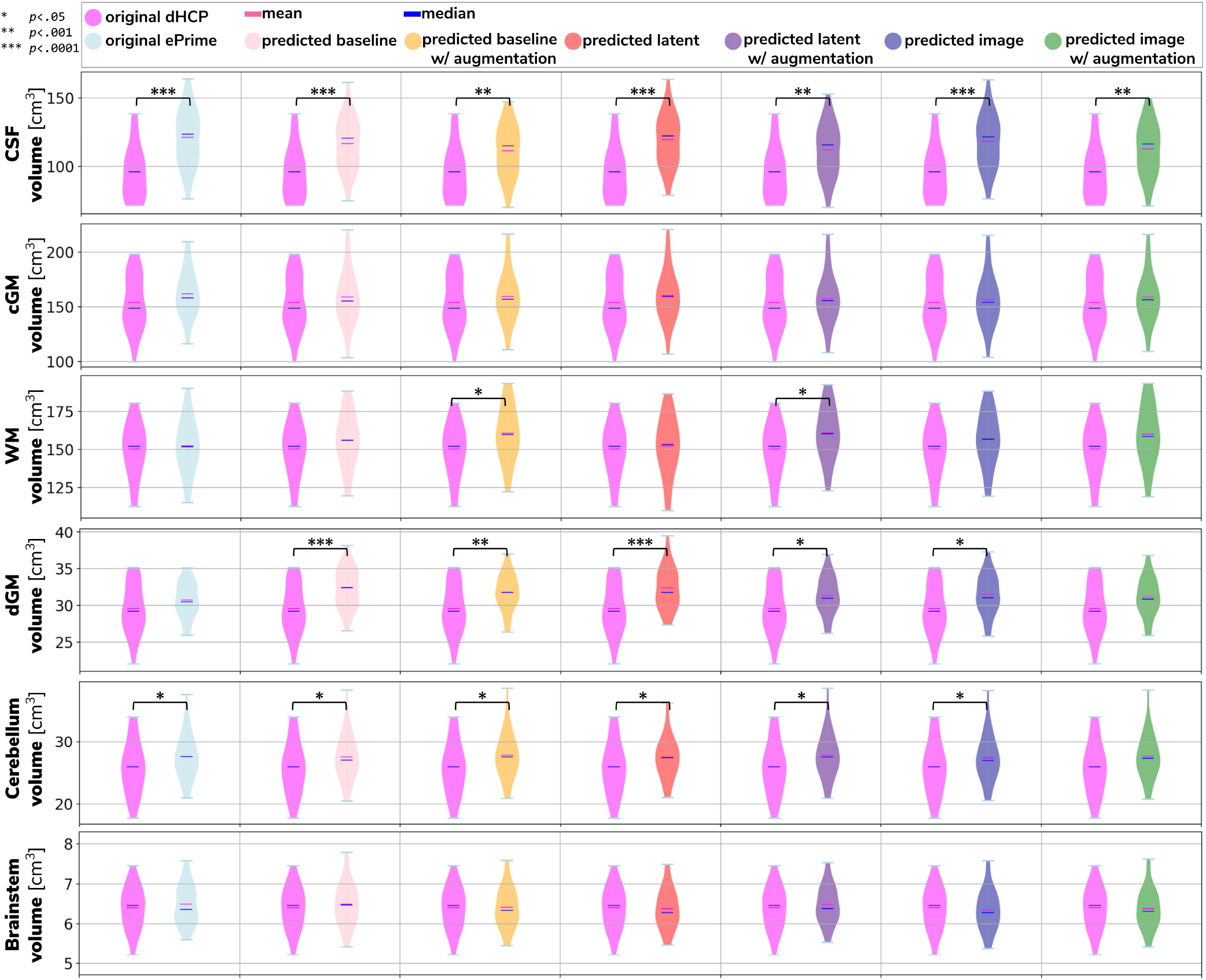
Volume measures of CSF, cGM, WM, dGM, cerebellum and brainstem in our test datasets. In magenta we show the original dHCP tissue volumes, while in light blue we show the original ePrime tissue volumes. The results of the linear regression are reported above groups which showed significant differences in terms of cohort.

#### Cortical thickness

Figure 7 summarizes the results of applying the cortical thickness algorithm on the predicted segmentation maps for all six methods. Before harmonisation, the matched subsets from the dHCP and ePrime cohorts showed a significant difference in mean cortical thickness (dHCP: *M* = 1.73, *SD* = 0.12; ePrime: *M* = 1.93, *SD* = 0.13; *t*(58) = 6.33, *p* < .001). After applying the harmonisation to the ePrime sample, mean cortical thickness no longer differed between the two subsamples for four of our methods. These results are summarised in panel H from Figure 7, where the models which obtained harmonised values in terms of mean cortical thickness measures are shown in bold. Figure 7 also shows the association between PMA and mean cortical thickness before (panel A) and after applying the models (panels B-G) on the matched dHCP and ePrime subsets. A linear model regressing unharmonised mean cortical thickness on PMA, GA, sex, and cohort revealed a significant effect of cohort (*β* = 0.20; *p* < .001), consistent with a group difference in mean cortical thickness reported above, as well as a significant effect of PMA (*β* = 0.04; *p* < .001), consistent with an increase in cortical thickness with increasing PMA. After applying the methods, the effect of cohort was rendered non-significant for four of the methods (see highlighted panels C, E, F, G from Figure 7), while the effect of PMA remained stable across all six methods.

**Figure 7.**
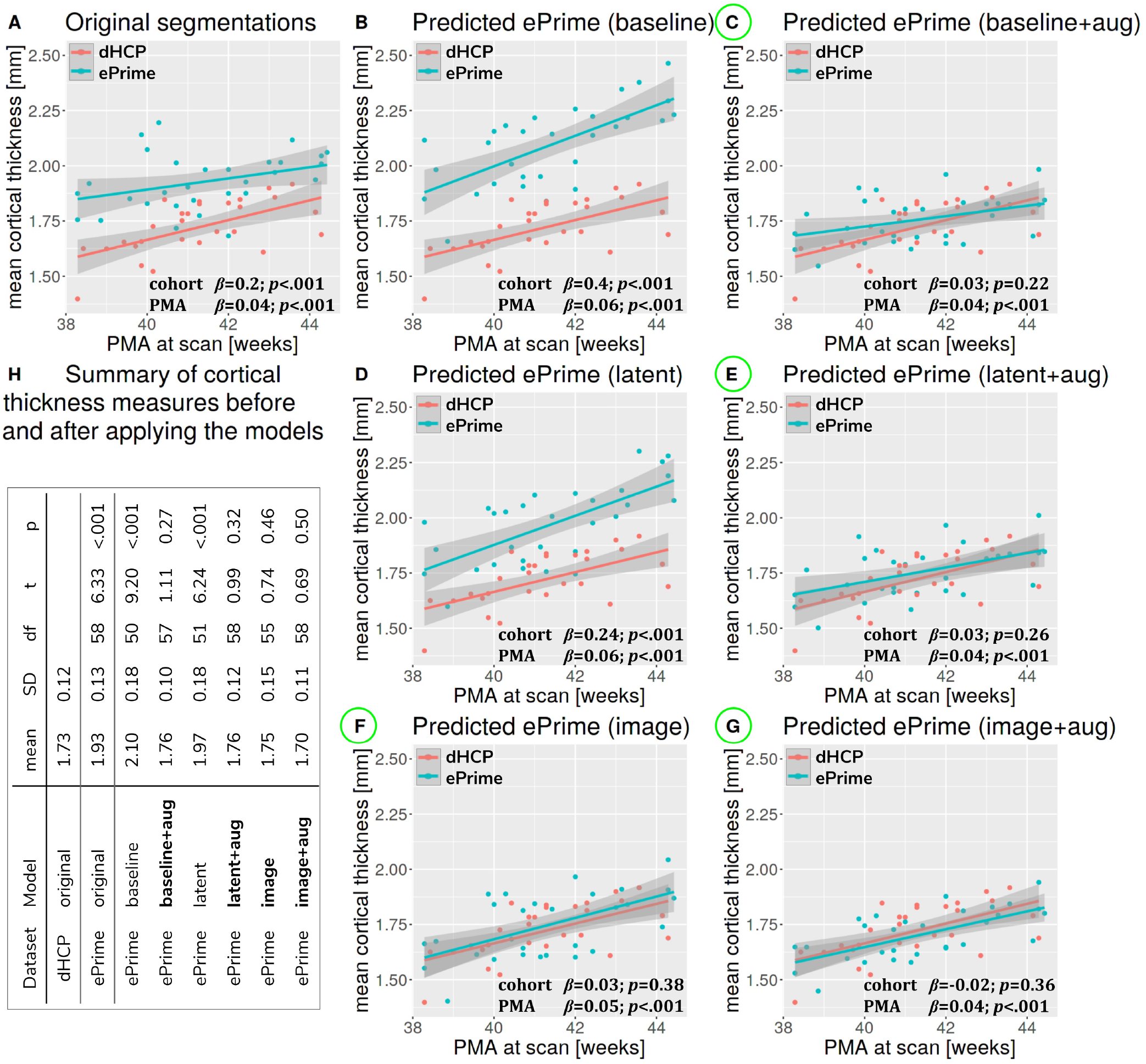
Mean cortical thickness measures computed for the two dHCP and ePrime subsamples with similar GA and PMA.

We performed a similar analysis on thickness measures of the cortical substructures. To obtain these measures, we used the original and the predicted cortical gray matter segmentation maps (obtained by applying each of our six methods) as input to the trained cortical parcellation network to predict cortical substructure masks. We then used these masks to calculate local cortical thickness measures. Our results are summarised in Figure 8.

**Figure 8.**
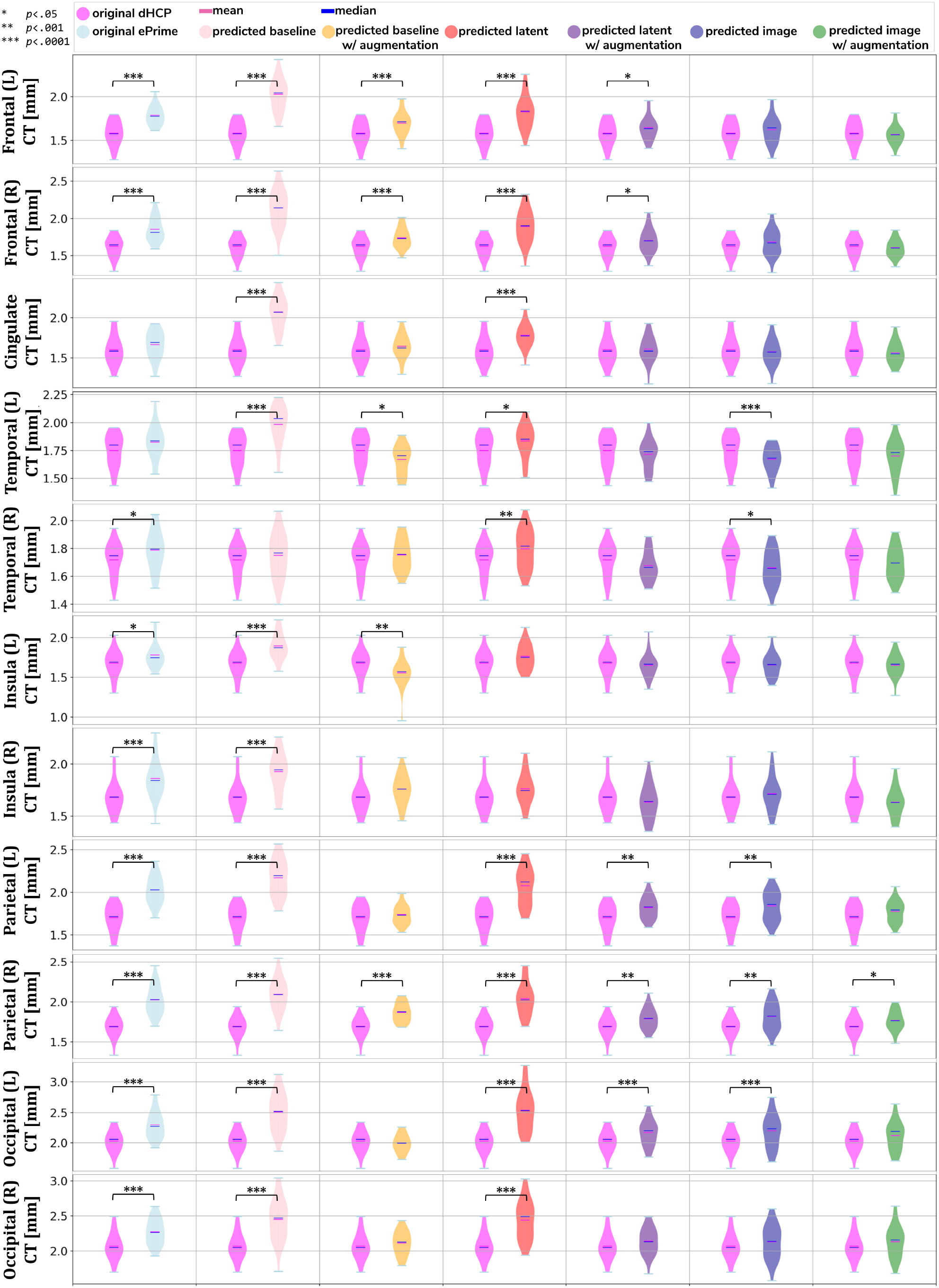
Local mean cortical thickness measures before (first column) and after (columns 2-7) applying the models. The results of the linear regression are reported above groups which showed significant differences in terms of cohort.

#### Example Predictions

To further narrow down which of the four remaining methods was best at harmonising our ePrime neonatal dataset, we looked at the predicted segmentations. Figure 9 shows two example neonates from the ePrime dataset with GA = 32.9w, PMA = 43.6w, and with GA = 28.7w, PMA = 44.7w, respecitvely. The first column shows *T*_2_w saggittal and axial slices, respectively, while the following four columns show example tissue prediction maps produced by the four models: *baseline with augmentation, latent with augmentation, image* and *image with augmentation*, respectively. Although all four methods performed well in terms of harmonising tissue segmentation volumes and global mean cortical thickness values for the two subsamples with similar GA and PMA, previously presented quantitative results as well as the example above suggest that the *image with augmentation* method was more robust.

**Figure 9.**
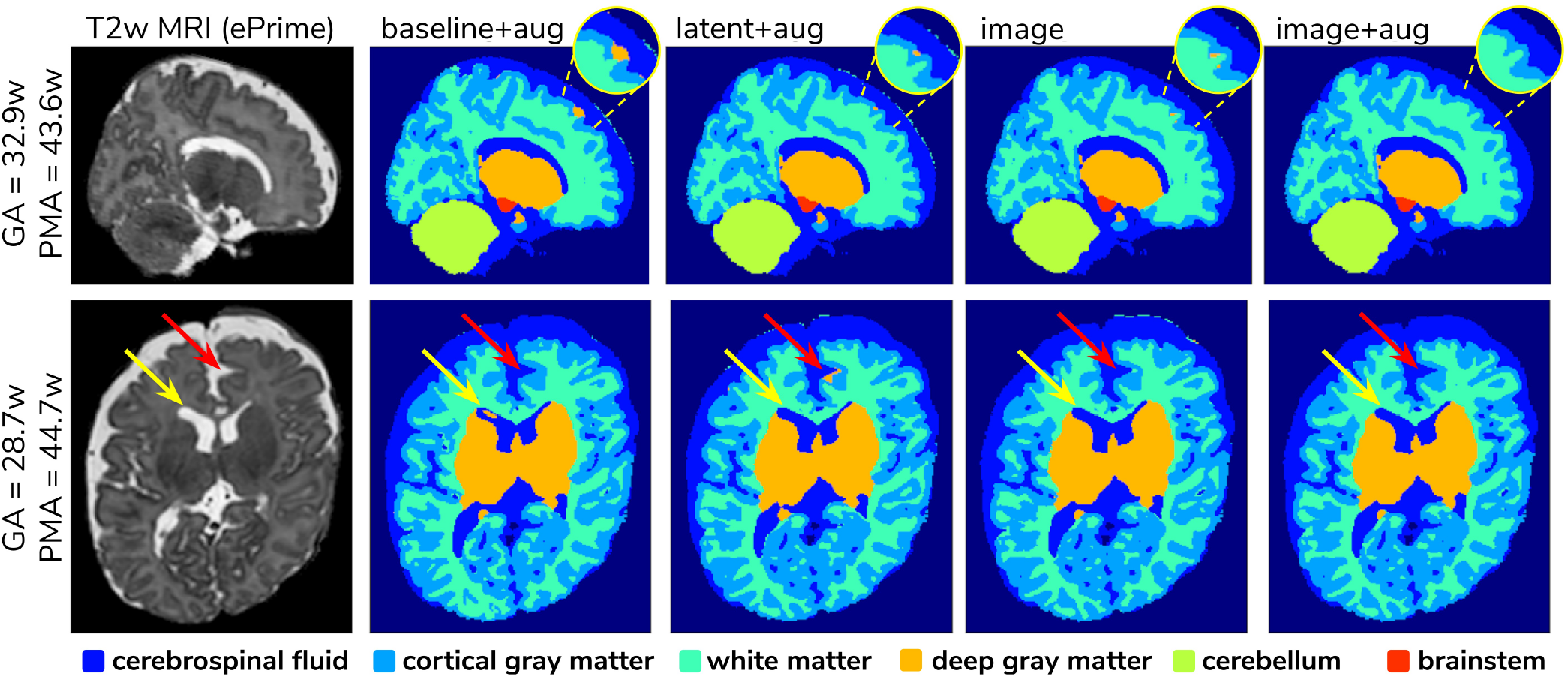
Example predicted segmentation maps for the best performing models. On the first row we show an example where three of the models (*baseline with augmentation, latent with augmentation* and *image*) misclassified a part of the cortex as being deep gray matter. This is more pronounced in the *baseline with augmentation* model, while the *latent with augmentation* and *image* show a slight improvement. The *image with augmentation* model corrected the problem entirely. On the second row the yellow arrow points to an area of CSF where the *baseline with augmentation* model misclassified it as dGM, while the other three models did not have this problem. The red arrow on the other hand points to an area where the *latent with augmentation* model misclassified cGM as deep gray matter. This problem does not appear in the other models.

#### 3.3 Analysis of harmonised cortical substructures

In this section we analyze the harmonised cortical gray matter segmentation maps using the *image with augmentation* model. We produce tissue segmentation maps for the entire ePrime dataset and calculate cortical thickness measures on the predicted and ground truth cortical gray matter tissue maps of both cohorts. In addition, we use the trained cortical parcellation network to produce cortical substructure masks. We perform a term *vs* preterm analysis on the harmonised cortical gray matter maps and we show the importance of harmonising the data with a proof-of-principle application setting where we investigate the association between cortical thickness and a language outcome measure.

##### Comparison of term and preterm cortical maps

Associations between cortical thickness and GA or PMA in the full dHCP and ePrime datasets (excluding subjects with PMA > 45 weeks and PMA < 37 weeks at time of scan) for the whole cortex are depicted in Figure 10, where we show individual regression lines for preterm-born and term-born neonates. The first column consists of dHCP-only subjects, while the following two columns showcase both cohorts together, before and after harmonising the cortical gray matter tissue maps.

**Figure 10.**
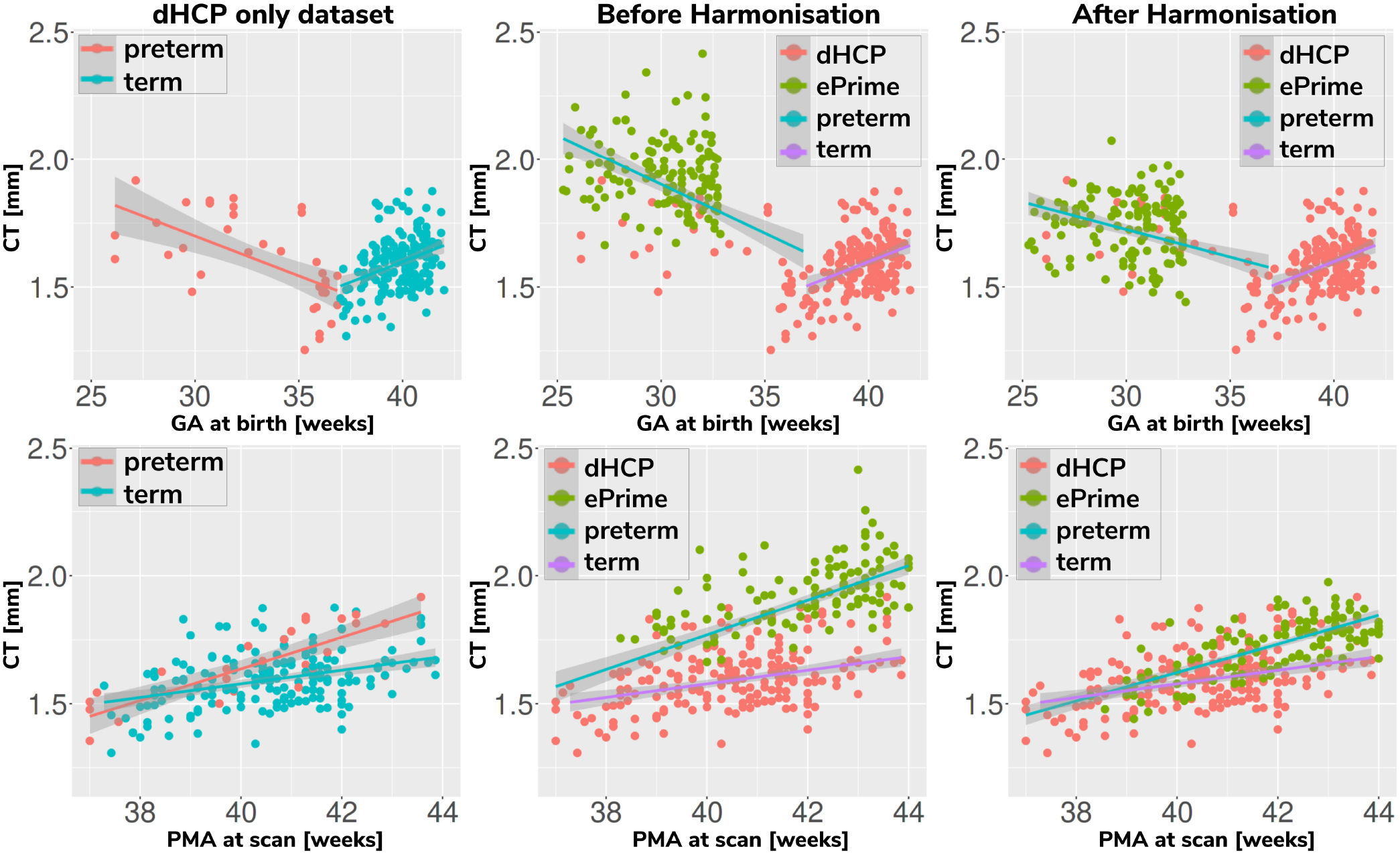
Mean cortical thickness measures in our dHCP dataset (first column), and in both of our cohorts before (second column) and after (third column) harmonising the tissue segmentation maps. The first row plots the cortical thickness measures against GA, while the second row plots the cortical thickness measures against PMA, with individual regression lines on top.

A linear model regressing dHCP-only mean cortical thickness on PMA, GA, sex, birth weight and the interaction between PMA and GA revealed a significant effect of PMA (*β* = 0.19; *p* < 0.001), a significant effect of GA (*β* = 0.16; *p* = 0.002), and a significant effect of the interaction between PMA and GA (*β* = –0.004; *p* = 0.002), indicating that infants born at a lower GA showed a stronger relationship between PMA and CT. When performing the same analysis in the pooled ePrime and dHCP data before harmonising the maps, the effect of GA and the effect of the interaction were rendered not significant (GA: β = 0.009; *p* = 0.7 and PMA*GA: β = –0.0006; *p* = 0.5, respectively). This is corrected after harmonising the tissue maps, where the effects of GA (*β* = 0.06; *p* = 0.02) and the effects of the GA and PMA interaction (*β* = –0.001; *p* = 0.02) are, again, significant.

The second and third columns of Figure 10 show that after harmonising the tissue segmentation maps, the ePrime preterm-born neonates (green dots) are brought downwards into a comparable range of values to the dHCP preterms (red dots). Moreover, when plotting the cortical thickness measures against PMA, after harmonising the tissue maps, the intersection between the two individual regression lines (term and preterm-born neonates) happens at roughly the same age (PMA = 38.5 weeks) as in the dHCP-only dataset.

We extended the term *vs* preterm analysis on cortical thickness substructures. Figure 11 shows the results of applying a linear model regressing mean cortical thickness measures on PMA, GA, sex, birth weight and prematurity, where significant differences (*p* < 0.05) between the two cohorts (term and preterm-born neonates) are highlighted in the image.

**Figure 11.**
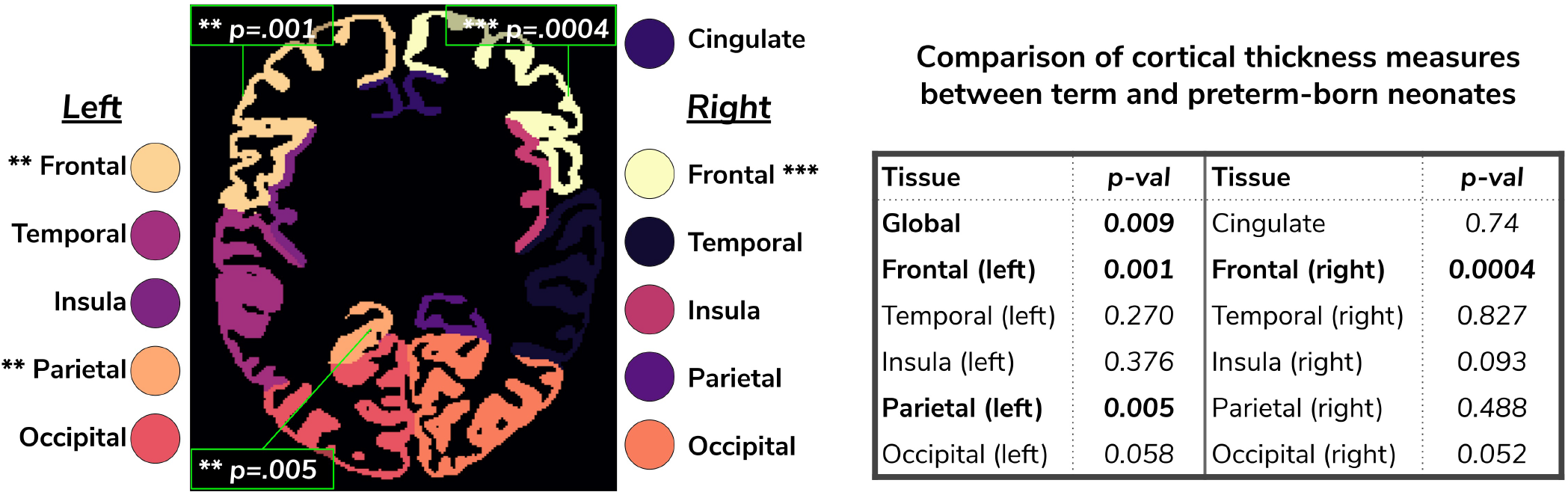
Comparison of cortical thickness measures for the whole cortex and for each of the 11 cortical subregions between term and preterm-born neonates. The results of the linear regression are reported in the table in terms of differences between term and preterm-born neonates.

##### Behavioural outcome association

As a final proof-of-principle, we demonstrate the importance of data harmonisation in an application setting investigating the association between neonatal cortical thickness and a behavioural outcome measure. For this, we consider language abilities as assessed between 18 and 24 months in both dHCP and ePrime cohorts using the Bayley Scales of Infant and Toddler Development (Bayley, 2006). Age-normed composite language scores were available for 203 toddlers from the dHCP cohort (M = 96.43; SD = 14.89) and 136 toddlers from the ePrime cohort (M = 91.25; SD = 17.37). For the neonatal cortical thickness measure, we focus on the left and right frontal cortex for illustration.

Regressing composite language score against ground truth left or right frontal cortical thickness in each cohort separately, controlling for PMA, GA, sex and intracranial volume showed that there was no significant association between neonatal left/right frontal cortical thickness and language abilities at toddler age in either of the cohorts. However, when pooling data from both cohorts together and rerunning the same analysis (using un-harmonised, ground truth cortical thickness), a significant association between left/right frontal cortical thickness and language abilities is seen (left: *β* = –17.56, *p* < 0.05, right: *β* = –18.76, *p* < 0.05), suggesting that greater frontal cortical thickness at term-equivalent age is associated with reduced language abilities at toddler age.

However, as can be seen in Figure 12, this is likely a spurious effect due to (artefactually) heightened cortical thickness values in un-harmonised ePrime data combined with lower language composite scores in the ePrime cohort (consistent with effects typically observed in preterm cohorts). Indeed, when rerunning the same analysis on harmonised data pooled across both cohorts, the effect of cortical thickness on language ability is rendered non-significant in both left (*β* = –13.99, *p* = 0.15) and right (*β* = –16.69, *p* = 0.068) frontal cortex, consistent with the ground-truth findings in each individual cohort.

**Figure 12.**
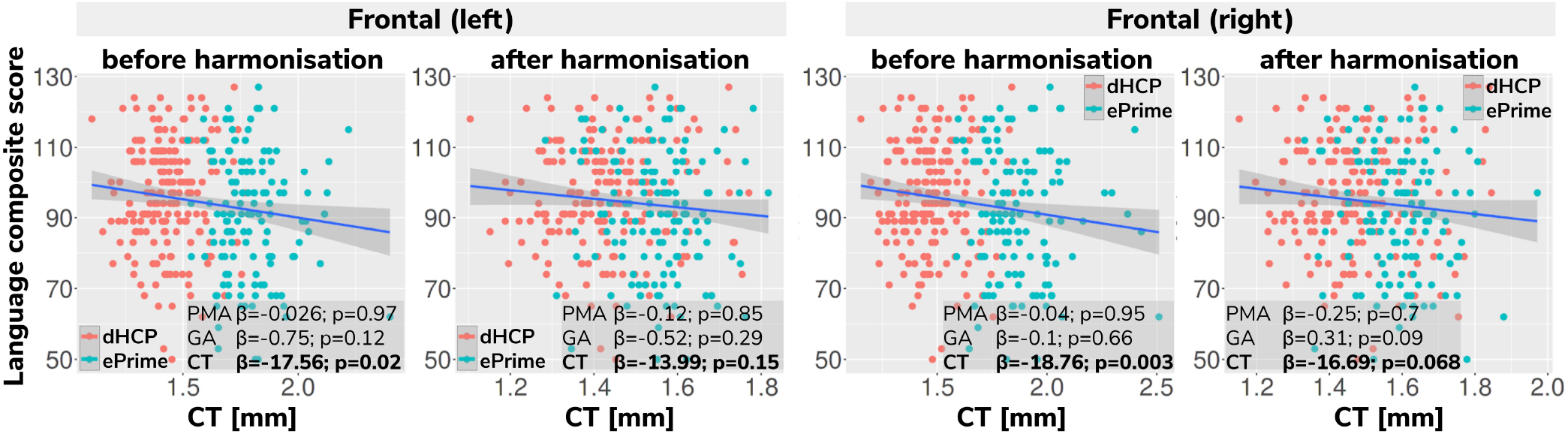
Language composite score against predicted left and right frontal cortical thickness measures before and after harmonising the tissue segmentation maps

## 4 DISCUSSION AND FUTURE WORK

In this paper we studied the application and viability of unsupervised domain adaptation methods for harmonising tissue segmentation maps of two neonatal datasets (dHCP and ePrime). We proposed an image-based domain adaptation model where a tissue segmentation network is trained with real dHCP and fake ePrime-like *T*_2_w 3D MRI volumes. The generator network was trained to produce realistic images in order to fool a domain discriminator, while also minimizing an NCC loss which aimed to enforce image similarity between real and synthesised images (Grigorescu et al., 2020). We trained this model using dHCP ground truth segmentation maps, and we compared it with a baseline 3D U-Net (Ronneberger et al., 2015), and a latent space domain adaptation method (Kamnitsas et al., 2017). The three methods were trained with and without data augmentation (Pérez-García et al., 2020).

We then analysed the extent to which each of the 6 trained models managed to harmonise the tissue segmentation maps of our two cohorts, by looking at tissue volumes and mean cortical thickness measures between subsamples of the dHCP and ePrime cohorts which showed comparable GA at birth and PMA at time of scan. Our results showed that our proposed model (*image with augmentation*) harmonised the predicted tissue segmentation maps in terms of cortical gray matter, white matter, deep gray matter, cerebellum and brainstem volumes (Figure 6). In terms of mean global cortical thickness measures, four of the trained methods (*baseline with augmentation*, *latent with augmentation*, *image* and *image with augmentation*) achieved comparable values when compared to the dHCP subset. In fact, we hypothesize that these four methods provided the best overall results because either they were trained using data augmentation or they acted as a deep learning-based augmentation technique (Sandfort et al., 2019), which made the segmentation network more robust to the different contrast and acquisition protocol of the ePrime dataset.

Using the cortical parcellation network, we also produced cortical thickness measures for the 11 cortical subregions (see Table 2). Again, the models trained with augmentation performed better than their no augmentation counterparts (see Figure 8). However, our proposed *image with augmentation* model performed best, whereby ePrime values, tending towards higher values before harmonisation, were brought downwards into a comparable range of values to dHCP, for 10 out of 11 cortical subregions (see Figure 8 last column). For the right parietal lobe, our proposed method outperformed the original segmentations and the other 5 models, but did not manage to bring the values down to a non-significant range. One potential reason for this is that, on a visual insepction, the ePrime cohort appears to suffer from more partial volume artifacts than its dHCP counterpart, which can confuse the segmentation network and can lead to overestimation of the cortical gray matter / cerebrospinal fluid boundary. Moreover, a close inspection of the predicted tissue segmentation maps (see Figure 9) also showed that our proposed model (*image with augmentation*) corrected misclassified voxels which were prevalent in the other 3 methods.

We used the harmonised cortical segmentation maps to look at differences in both global and local cortical thickness measures between term and preterm-born neonates. We showed in Figure 11 that our harmonised cortical gray matter maps resulted in global thickness measures which were comparable with the dHCP-only neonates, while also revealing a significant effect of GA and the interaction between age at scan and at birth. We performed a similar analysis on the local cortical thickness measures and highlighted three regions of interest (frontal left, frontal right, and parietal left) which showed significant differences between the two cohorts (see Figure 11). These regions are consistent with previous studies (Nagy et al., 2011) where cortical thickness measures were shown to differ in preterm-born neonates when compared to term-born neonates in an adolescent cohort.

Finally, we showed the importance of harmonising the cortical tissue maps by investigating the association between neonatal cortical thickness and a language outcome measure. After harmonisation, regressing language composite score against predicted left or right frontal cortical thickness in the two pooled datasets, showed no significant effect of cortical thickness (second column of Figure 12), consistent with the ground-truth results seen in each cohort individually. This analysis demonstrates that without data harmonisation, pooling images from separate datasets can lead to spurious findings that are driven by systematic differences in acquisitions rather than by true underlying effects. Our harmonisation allows for our two datasets to be combined into joint analyses while preserving the underlying structure of associations with real-world outcomes.

Our study was focused on unsupervised domain adaptation approaches; in future we would like to investigate semi-supervised approaches as well by including reliable segmentations of the ePrime cohort. Moreover, the latent based domain adaptation method was trained using the features at each layer of the decoding branch, without analysing different combinations of the encoding-decoding layers. In future, we aim to extend our work to harmonise diffusion datasets.

## AUTHOR CONTRIBUTIONS

I. G. prepared the manuscript, implemented the code for the domain adaptation models and the analysis. L.V. participated in the implementation of the analysis code, the study design and interpretation of the results. A.U. assisted with data preprocessing, design of the study and interpretation of the results. D.B. performed preprocessing of the dHCP and ePrime datasets. L.C.-G. developed MRI acquisition protocols for the neonatal dHCP datasets. C.N. participated in the study design and interpretation of the results. A.D.E., J.V.H. are coordinators of the dHCP project. M.M. supervised all stages of the current research. M.D. conceptualised the study, supervised all stages of the current research and preparation of the manuscript. All authors gave final approval for publication and agree to be held accountable for the work performed therein.

## ACKNOWLEDGMENTS

We thank everyone who was involved in acquisition and analysis of the datasets. We thank all participants and their families. The views expressed are those of the authors and not necessarily those of the NHS, the NIHR or the Department of Health. This paper is an extension of our previous work (Grigorescu et al., 2020).

## SUPPLEMENTAL DATA

### DATA AVAILABILITY STATEMENT

The dHCP datasets analyzed for this study will become available after the public release of the dHCP data. The code developed for this study will become available online after publication of the article.

## Footnotes

1 http://www.developingconnectome.org/

2 https://www.npeu.ox.ac.uk/prumhc/eprime-mr-imaging-177

